# Somatic mutations in CTCF zinc fingers produce cellular phenotypes explained by structure-function relationships

**DOI:** 10.1101/2021.01.08.425848

**Authors:** Charles G Bailey, Shailendra Gupta, Cynthia Metierre, Punkaja MS Amarasekera, Patrick O’Young, Wunna Kyaw, Tatyana Laletin, Habib Francis, Crystal Semaan, Krishna P Singh, Charles G Mullighan, Olaf Wolkenhauer, Ulf Schmitz, John EJ Rasko

## Abstract

**Background:** Human cancers commonly contain mutations in transcription factors that lead to aberrant DNA binding or altered effector function at target sites. One such factor significantly mutated in cancer is the evolutionarily-conserved CCCTC-binding factor (CTCF), which has fundamental roles in maintaining chromatin architecture and transcriptional regulation. Numerous cancer genome sequencing and functional studies have revealed CTCF’s role as a haploinsufficient tumour suppressor gene. However, to date, structure-function relationships of somatic CTCF mutations have not been examined.

**Methods:** We collated somatic CTCF mutations from cancer genome portals and published studies to determine their nature, frequency, distribution and potential functional impact. We undertook an in-depth examination of 5 CTCF missense zinc finger (ZF) mutations occurring within key intra- and inter-ZF residues. We performed functional analyses including cell growth, colony-formation, chromatin immunoprecipitation and transcriptional reporter assays. Based on their homology, each ZF mutation was then modelled on CTCF’s ZF domain crystal structure and its structural impact analysed using molecular dynamics simulations.

**Results:** We observed an enrichment of somatic missense mutations occurring in the ZF region of CTCF, compared to the unstructured N- and C-termini. Functional characterisation of CTCF ZF mutations revealed a complete (L309P, R339W, R377H) or intermediate (R339Q) abrogation as well as an enhancement (G420D) of the anti-proliferative effects of CTCF. DNA binding at select sites was disrupted and transcriptional regulatory activities abrogated. *In silico* mutagenesis revealed that L309P had the highest mutation energy and thus most severe impact on protein stability. Molecular docking and molecular dynamics simulations confirmed that mutations in residues specifically contacting DNA bases or backbone exhibited loss of DNA binding (R339Q, R339W, R377H). Remarkably, R339Q and G420D were stabilised by the formation of new primary DNA bonds. All mutations exhibited some loss or gain of bonds at neighbouring residues, often in adjacent zinc fingers.

**Conclusions:** Our data confirm the significant negative impact haploinsufficient CTCF ZF mutations have on its tumour suppressor function. A spectrum of loss-, change- and gain-of-function impacts in CTCF zinc fingers are observed in cell growth regulation and gene regulatory activities. We have established that diverse cellular phenotypes in CTCF are explained by examining structure-function relationships.

## Background

Comprehensive catalogues of somatic mutations have been assembled from surveying the genomic landscape in numerous human cancers. More than 200 large-scale studies involving cancer types or subtypes of clinical or societal importance have been deposited in the cBio Cancer Genome Portal^1^. These studies have provided new insights into cancer causation and offered new leads for potential therapeutic intervention using a genomics-driven oncology approach. Cancers are remarkably heterogeneous in their distribution and frequency of somatic mutations. Paediatric cancers contain as few as 0.1 mutations per megabase (Mb), whereas lung and melanoma samples may accumulate over 100/Mb (average 4.0/Mb)^2^. Whilst some genes are mutated at high frequencies, most genes are mutated at intermediate frequencies (2-20%)^3^ adding to the complex molecular landscape underlying tumour biology. Those genes that exhibit mutation frequencies above background have been called significantly mutated genes, of which 127 have been identified amongst a dozen cancers^4^. These mutations disrupt diverse cellular processes including transcriptional regulation, histone modification, genome integrity, signalling and splicing^4^. Mut-driver genes (of which 138 have been identified) are a similar concept, whereby mutation or inactivation can cause a selective growth advantage in a direct or indirect manner^5^. Examples include inactivation of a tumour suppressor gene or activation of oncogenes.

In tumour cells, recurrent acquired mutations have been observed in nearly every DNA, RNA and protein component of normal transcriptional control^6^. These somatic mutations may directly impact transcription factors (TFs), their target sites, *cis*- and *trans*-regulatory elements as well as chromatin architecture leading to transcriptional dysregulation in cancer. Dysregulation of transcriptional programs in cancer cells can lead to transcriptional dependencies that offer opportunities for exploitation with targeted therapeutic strategies^6^. For example, pharmacological inhibition of the BET bromodomain-containing BRD4 protein has emerged as a promising therapeutic strategy to prevent MYC-dependent transcriptional signaling in various haemopoietic malignancies^7–10^. Investigating and exploiting these acquired cellular vulnerabilities is a major thrust of many cancer research efforts.

Approximately 1,600 to 2,000 TFs have been validated or predicted within the human genome^11,12^. TFs containing the zinc-coordinating C2H2 class of DNA binding domains represent the largest class of transcription factors^11^, comprising nearly 50% of all TFs^13^. Human C2H2 TFs contain an average of ~10 ZFs, specifying target sites of ~30 bases^14^, however not all ZFs contact DNA simultaneously or indeed, are involved in DNA binding. Furthermore, the impact of somatic mutations on many TFs is unknown. Nor is it known whether such changes impact DNA binding or transcriptional activation globally or in a locus-specific manner.

One such C2H2-ZF-containing transcription factor, CCCTC-binding factor (CTCF), features a tandem array of 11 ZFs enabling multivalent binding to DNA target sites. Careful mutational analysis of key residues co-ordinating Zn^2+^ ion binding and ZF formation have shown key central ZFs that contribute binding to a core consensus site, whilst peripheral ZFs stabilise CTCF binding and bind additional conserved and non-conserved motifs^15^. Through combinatorial DNA binding and ZF multivalency, CTCF plays diverse roles in transcriptional regulation and three-dimensional genome organisation such that it has been called the ‘master weaver’ protein^16^. Unprecedented insights into nuclear organisation, obtained from high-resolution conformational maps of chromatin interactions, have defined the rules governing CTCF-mediated chromatin organisation. Firstly, CTCF links gene regulation to genomic architecture by co-ordinating DNA looping together with cohesin^17–19^. Secondly, CTCF defines the boundaries of topologically associating domains (TADs)^20–22^ in a structural framework that is evolutionarily conserved^23^. Depletion of CTCF can result in loss of DNA looping and insulation within TADs, however genomic compartmentalisation is preserved^24^. Lastly, TAD organisation is CTCF site orientation-specific^23,25^, such that rewiring or inverting CTCF sites can significantly perturb gene expression by affecting promoter-enhancer interactions or boundaries between euchromatin and heterochromatin^26–28^.

CTCF plays an integral role in cell-type specific genomic organisation and development. CTCF’s role in development and differentiation has been examined in at least seven tissues or developmental stages in mice, zebrafish and Drosophila^29^. CTCF is absolutely essential, as CTCF null embryos are unable to implant^30^ and maintenance of CTCF expression ensures somatic cell viability^31^. Extensive characterisation of the action of CTCF *in vitro* and *in vivo* has led to its classification as a haploinsufficient tumour suppressor gene^31–33^. Whilst isolated somatic CTCF mutations were first identified in some solid tumours^34^, numerous cancer genome studies since have highlighted the impact and prevalence of CTCF mutation in multiple cancers^4^. CTCF is a significantly mutated gene in ~20% of endometrial cancers^35–38^ and is recurrently mutated in myeloid and lymphoid malignancies^39–42^.

Despite many CTCF mutations having been identified in numerous cancer types, the functional consequences of these mutations have not been thoroughly examined. In this study, we performed a meta-analysis of all publicly available cancer mutation data for CTCF and showed a significant enrichment of missense mutations occurring in CTCF’s ZF DNA binding domain. We have functionally characterised a subset of representative ZF mutations detected in acute lymphoblastic leukaemia samples to examine their consequences. Finally, we compared the impact of CTCF ZF mutation on DNA binding, transcriptional activation as well as on CTCF ZF domain structure using molecular modelling and molecular dynamics simulations. This is the first study to examine the effect of somatic mutation on CTCF ZF structure-function relationships.

## Results

### CTCF ZF domain is enriched for somatic missense mutations in cancer

We analysed cancer genome sequencing databases and published mutation data to determine the distribution, frequency and nature of somatic mutations occurring in CTCF (Supplementary Table 1). The distribution and frequency of all known somatic mutations in CTCF is shown with recurrent mutant residues indicated (Figure 1A). The recurrent T204fs*26 and T204fs*18 mutations in CTCF arise due to a high frequency of insertions or deletions within a 30 bp purine-rich (>85%) region at c.1048 –c.1077 encoding T204. Frequently occurring missense or nonsense mutations occur at H284, S354, R377, R448 and R457 within the ZF region of CTCF (Figure 1A). Further analysis revealed that inactivating nonsense and frameshift mutations account for ~40% of somatic CTCF mutations (Figure 1B). This result exceeds the ‘20/20 rule’ for tumour suppressor gene classification which requires that >20% of somatic mutations are inactivating^5^ and affirms our earlier work demonstrating CTCF’s role as a tumour suppressor^31–33^. CTCF mutations occur prominently within hormone-responsive cancers arising in the endometrium and breast (~48%) (Figure 1C).

**Figure 1.**
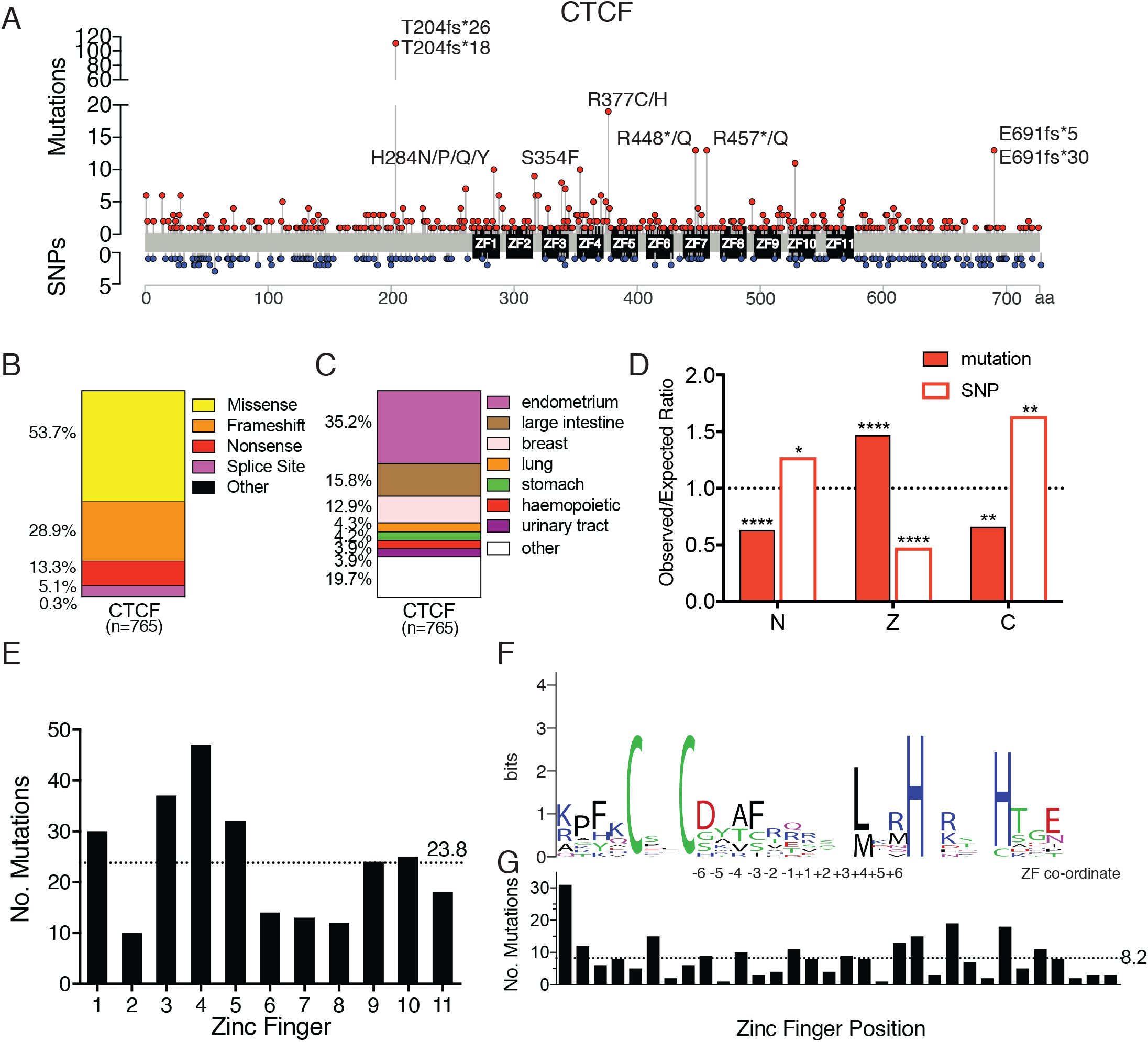
Distribution and impact of CTCF somatic mutations in cancer. **(A)** The landscape of somatic mutations (above) and SNPs (below) occurring in CTCF: the distribution and frequency within the coding region is shown, recurrent somatic mutations (occurring ≥10 times) are labelled. For a curated list of non-redundant CTCF mutations from cancer genome sequencing studies (TCGA, COSMIC) and published studies see Supplementary Table 1. *CTCF* mutation type **(B)**; and tissue distribution **(C)** are shown; n=total number of mutations. **(D)** Analysis of cancer-related somatic missense variants and missense SNPs occurring in each domain of CTCF. The expected occurrence was calculated from the total number with the proportion of missense variants expected in each domain if they were evenly distributed. The observed/expected ratio confirms if there is a de-enrichment (<1.0) or an enrichment (>1.0) of non-synonymous changes. **(E)** Frequency of somatic missense mutations occurring in specific ZFs of CTCF, the mean for all ZFs is shown (dotted line). **(F)** Sequence logo of all 11 aligned CTCF ZFs; numbers (−6 to +6) indicate co-ordinates within the DNA-binding portion of the ZF. Similar amino acids are coloured: black – hydrophobic (G, A, V, I, L, P, W, F, M); green – polar (S, T, Y, C); purple – polar amide (Q, N); blue – basic (K, R, H); and red – acidic (D, E). The height of each amino acid residue is proportional to its observed frequency. The overall height of each letter ‘stack’ is proportional to the sequence conservation, shown in bits **(G)** Frequency of missense somatic mutations at each ZF position; the mean for all ZFs is shown (dotted line). Data represent the mean ± SD with statistical analysis performed using the Chi-square test (*, p<0.05; **, p<0.01; ****, p<0.0001).

We next examined somatic missense mutations and SNPs reported for CTCF and compared their observed and expected occurrences (Supplementary Table 2). CTCF’s ZF domain has a significant enrichment for somatic missense mutations observed over the number expected for its relative size, such that the observed/expected (O/E) ratio =1.47, (*p*<0.0001). Furthermore, there was a de-enrichment of non-synonymous SNPs occurring within the ZF domain (O/E=0.48, *p*<0.0001) (Figure 1D, Supplementary Table 2). These results suggest that the human CTCF ZF region is intolerant to normal genetic variation, but is frequently inactivated in cancer. As ZF mutations would likely affect DNA binding, these are likely to have a significant impact on CTCF function. There is a concomitant paucity of missense somatic mutations within the N- and C-termini of CTCF (O/E = 0.63, *p*<0.0001 and O/E = 0.65, *p* = 0.0269 respectively, Figure 1D, Supplementary Table 2). Strikingly, the opposite trend is observed for SNPs in CTCF with an enrichment of missense SNPs in the N-terminus (O/E = 1.27, *p* = 0.0269) and C-terminus (O/E=1.63, *p* = 0.0032) Figure 1D, Supplementary Table 2). We then determined the potential functional impact of somatic missense mutations in CTCF using Polyphen analysis. Missense mutations exhibited an overall greater functional impact than missense SNPs (0.80±0.35 vs 0.49±0.44, mean±SD, *p*<0.0001, Supplementary Figure 1A). Further analysis indicated that there was a decrease in the ratio of transition to transversion mutations when comparing SNPs to missense somatic mutations (2.24 to 1.19 respectively, *p*<0.0001, Supplementary Figure 1B). These data provide further support for the role of CTCF as a tumour suppressor that is frequently mutated and functionally impacted in cancer.

As the majority of somatic missense mutations in CTCF occur within the ZF domain we next analysed the distribution of missense mutations in specific ZFs of CTCF. We found that the greatest proportion of mutations occurred in ZF4 (~20%), followed by ZF3 (~15%) (Figure 1E). ZFs 3-7 have been shown to be responsible for binding CTCF’s core 15 bp consensus, with other ZFs providing binding specificity depending on adjacent motifs^15,43^. A sequence logo depicting all 11 ZFs in CTCF (10 C2H2- and 1 C2HC-type) shows the conserved Cys and His residues that co-ordinate Zn^2+^ binding, an invariant hydrophobic Leu or Met residue at +4 and substantial amino acid variation at other positions (Figure 1F). The proportion of mutations occurring at each position within ZFs was determined. This analysis revealed that the proportion of inter-ZF mutations was 31.5%, Cys/His mutations (17.7%) and those affecting key DNA binding residues (−1, +2, +3, +6, 15.6%). Thus, approximately one-third of missense CTCF ZF mutations have an unknown impact but likely affect ZF folding and stability.

### CTCF ZF mutations exhibit loss- and gain-of-function in cell growth phenotypes *in vitro*

To determine the functional consequences of CTCF ZF mutations, we examined missense mutations that had been detected in acute lymphoblastic leukaemia (ALL) samples: L309P (T-ALL; Mullighan unpublished) R339Q^39^, R377H^44^ and G420D (diagnosis and relapsed hyperdiploid B-ALL; Mullighan unpublished) (Figure 2A, Supplementary Table 1). R377H occurs within the inter-ZF region, L309P affects the conserved intra-ZF Leu/Met residue, whilst G420D and R339Q both occur at key DNA contacting residues +2 and +6, respectively (Figure 2A). We included R339W as a positive control as it was first identified in Wilms’ tumour as a potential ‘change-of-function’ mutation that abrogated DNA binding to a subset of CTCF sites regulating genes involved in cell proliferation^34^. All five mutations exhibit high Polyphen scores, indicating they significantly impact CTCF function (Figure 2A). We introduced these mutations into HA epitope-tagged human CTCF within a lentiviral expression vector that co-expresses eGFP via a 2A peptide^33^. We transduced K562 erythroleukemia cells with CTCF WT and mutant constructs and showed that ectopic CTCF expression occurred at similar levels and above endogenous CTCF levels (Figure 2B). Immunofluorescent staining for ectopic HA-tagged CTCF indicated that all CTCF mutants maintained nuclear localisation similar to WT CTCF (Figure 2C). We next examined cell growth and showed that WT CTCF overexpression suppressed cellular proliferation (*p*<0.0001) consistent with it being a tumour suppressor and as previously shown^33^ (Figure 2D). Mutants L309P, R377H and R339W abrogated the tumour suppressive effect of CTCF and exhibited cellular proliferation similar to the empty vector control (all *p*<0.0001 compared to WT), whilst R339Q had an intermediate effect on CTCF’s anti-proliferative function (*p*<0.0001 compared to WT; *p*<0.001 compared to control, Figure 2D). K562 cells expressing CTCF G420D exhibited similar proliferation to WT CTCF (Figure 2D). We next performed clonogenicity assays and showed that WT CTCF suppressed the colony-forming abilities of K562 cells as expected (*p*<0.0001, Figure 2E). Again, L309P, R377H and R339W abrogated the suppressive effect of CTCF on colony formation (*p*<0.0001) whilst R339Q had an intermediate effect compared to control (*p*<0.0001) and a near-intermediate effect compared to WT (*p* = 0.052). Remarkably, G420D exhibited gain-of-function by further reducing the clonogenic capacity compared to WT (Figure 2E).

**Figure 2.**
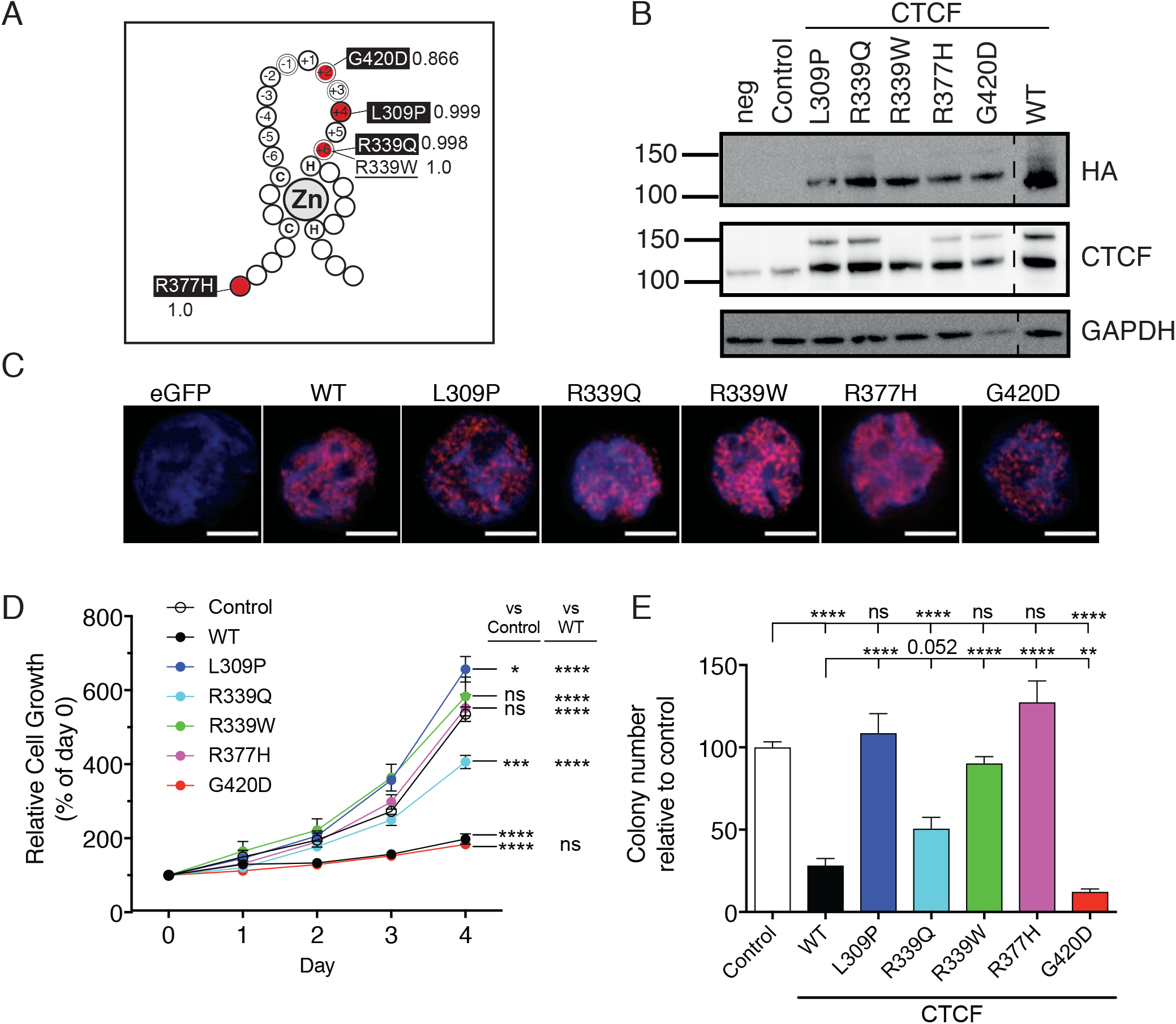
Functional impacts of CTCF ZF missense mutations. **(A)** Published and unpublished missense CTCF mutations (red circles) occurring in acute lymphoblastic leukaemia (L309P, R339Q, R337H, G420D - highlighted in black) superimposed on a C2H2 ZF structure: C = cysteine, H = histidine, Zn = Zn^2+^ ion; R339W (underlined) is a previously characterized change-of-function mutation used as a control. The Polyphen score for each mutation is indicated. Numbers (−6 - +6) indicate co-ordinates within the DNA-binding portion of the ZF; residues directly contacting DNA at positions −1, +2, +3 and +6 are indicated (white ring). **(B)** Western blot of WT and mutant CTCF expression in transduced K562 cells; anti-HA antibody detects ectopic CTCF, CTCF antibody detects total CTCF; GAPDH is a loading control; size markers (not shown) indicate MW in kDa; vertical dashed line indicates WT sample run on same blot but relocated for clarity. **(C)** Immunofluorescence of HA-tagged WT and mutant CTCF in K562 cells using anti-HA antibody, scale bar=5 μm. **(D-E)** Functional assays of CTCF mutants in K562 cells including: **(D)** MTT proliferation; and **(E)** colony forming assay in Methocult. Data represent the mean ± s.e.m for 3 experiments each performed in triplicate. Statistical analysis was performed using Mann-Whitney U-test (ns = not significant; *, *p*<0.05; **, *p*<0.01; ***, *p*<0.001; ****, *p*<0.0001).

### CTCF ZF mutations disrupt transcriptional activity

We next examined the impact of ZF mutations on transcriptional regulation by CTCF. Frequently occurring N-and C-terminal somatic missense mutations (Y226C and R603C respectively) were included as additional controls. Y226 is a key anchoring residue in the interaction of CTCF with the SA2-SCC1 cohesin complex^45^, whilst R603 resides within the nuclear localisation signal. Lentiviral plasmids encoding WT or mutant CTCF (Figure 3A) were transfected into HEK293T cells followed by quantitation of CTCF protein and eGFP fluorescence levels. All CTCF ZF mutants exhibited decreased levels of ectopic CTCF expression to levels ~10-20% of WT, whilst non-ZF mutants demonstrated levels comparable to, or higher than, WT control (Figure 3B & C). WT CTCF suppressed eGFP expression compared to eGFP control (*p*<0.0001, Figure 3C) whilst decreased eGFP expression was observed with CTCF mutants R339Q, R339W, R377H and G420D compared to WT. L309P had no impact; however, non-ZF mutations (Y226C and R603C) exhibited higher eGFP expression than WT (*p* = 0.0248 and *p* = 0.0061 respectively), but similar to empty vector control (Figure 3D). These data suggested that ectopic plasmid-encoded CTCF could regulate its own expression. This was supported by the prediction of over a dozen putative CTCF binding sites in the CTCF WT and mutant plasmid backbone including within the CMV promoter that drives viral RNA expression (horizontal dashes, Figure 3A). Accordingly, CTCF ZF mutants exhibited diminished CTCF protein expression and lower eGFP expression (Figure 3B, C & D). Collectively, these data indicate that CTCF ZF mutants impact on CTCF’s normal role as a transcriptional regulator, which most likely results from disruption or destabilisation of DNA binding.

**Figure 3.**
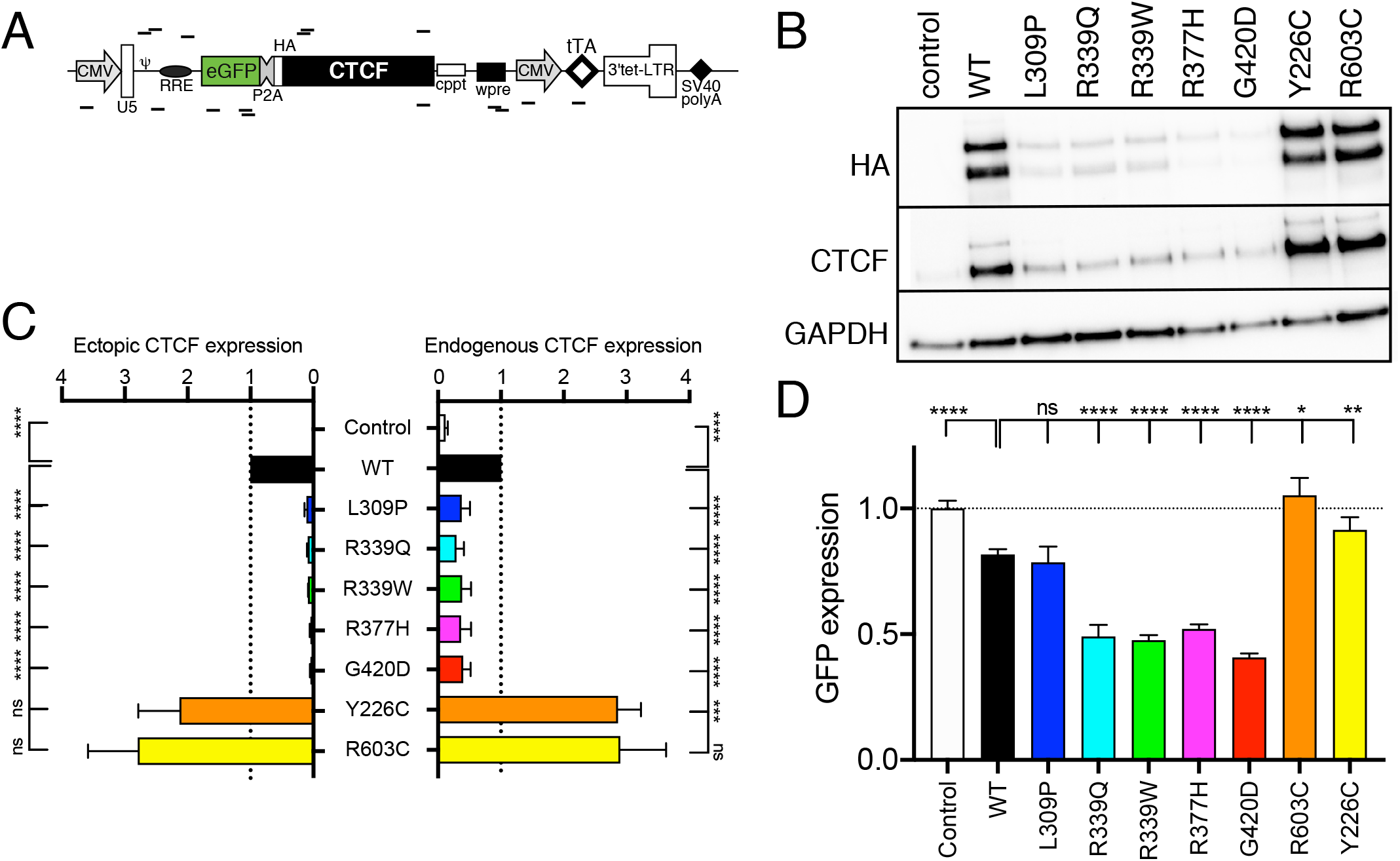
Differential gene regulatory activities exhibited by ZF-mutant CTCF. **(A)** Schematic of the lentivector plasmid used to measure WT and mutant CTCF gene regulatory activity. Predicted CTCF binding sites on both sense and antisense strands are represented by horizontal lines respectively. **(B-D)** Control (eGFP-alone), CTCF WT or mutant-containing lentivector plasmids were transfected into HEK293T cells for 48 h. **(B)** Representative Western blots (of 3 replicates) indicating ectopic (HA-tagged) CTCF, total CTCF and GAPDH loading control after transfection of HEK293T cells. **(C)** Densitometry of ectopic and endogenous CTCF expression normalised to GAPDH, shown relative to WT CTCF set as 1.0. **(D)** GFP mean fluorescence intensity (MFI) detected after 48 h and normalised to eGFP empty vector control set as 1.0. Data represent the mean ± s.e.m for 3 experiments each performed in triplicate except for the Western blots which are only single replicates. Statistical analysis was performed using Mann-Whitney U-test (ns = not significant; *, *p*<0.05; **, *p*<0.01; ***, *p*<0.001; ****, *p*<0.0001).

To examine this, we performed chromatin immunoprecipitation (ChIP) to determine if ZF-mutant disruption of transcriptional regulation leads to abrogation or alteration of DNA binding at CTCF target sites. Notably, we achieved equivalent levels of HA-tagged WT and ZF mutant CTCF in K562 cells after lentiviral transduction (~15-20% for all, Supplementary Figure 2). We then performed ChIP using an anti-HA antibody, followed by PCR amplification of known CTCF target sites (Figure 4). We observed both WT and mutant CTCFs still associating with archetypal CTCF target sites such as the *H19* imprinting control region (ICR) and the β-globin hypersensitivity site HS5 (Figure 4A). However, variegated CTCF mutant binding was detected at other cognate CTCF target sites proximal to the regulatory regions of *BAG1*, *MAGEA1*, *XIST*, *BRCA1*, *PLK* and *APPβ* (Figure 4A). All CTCF ZF mutants exhibited a selective loss of DNA binding, with L309P, R339Q and R337H mutations exhibiting the greatest loss in binding (Figure 4A-E). All CTCF mutants except G420D exhibited some loss of binding within the archetypal CTCF-regulated gene *C-MYC* (Figure 4B). CTCF binding sites within known enhancers (Figure 4C), insulator sites (Figure 4D) and TAD boundaries (Figure 4E) all showed selective binding by most CTCF ZF mutants. As CTCF binding is not completely abrogated, these data are consistent with CTCF ZF mutants displaying a change-of-function rather than loss-of-function.

**Figure 4.**
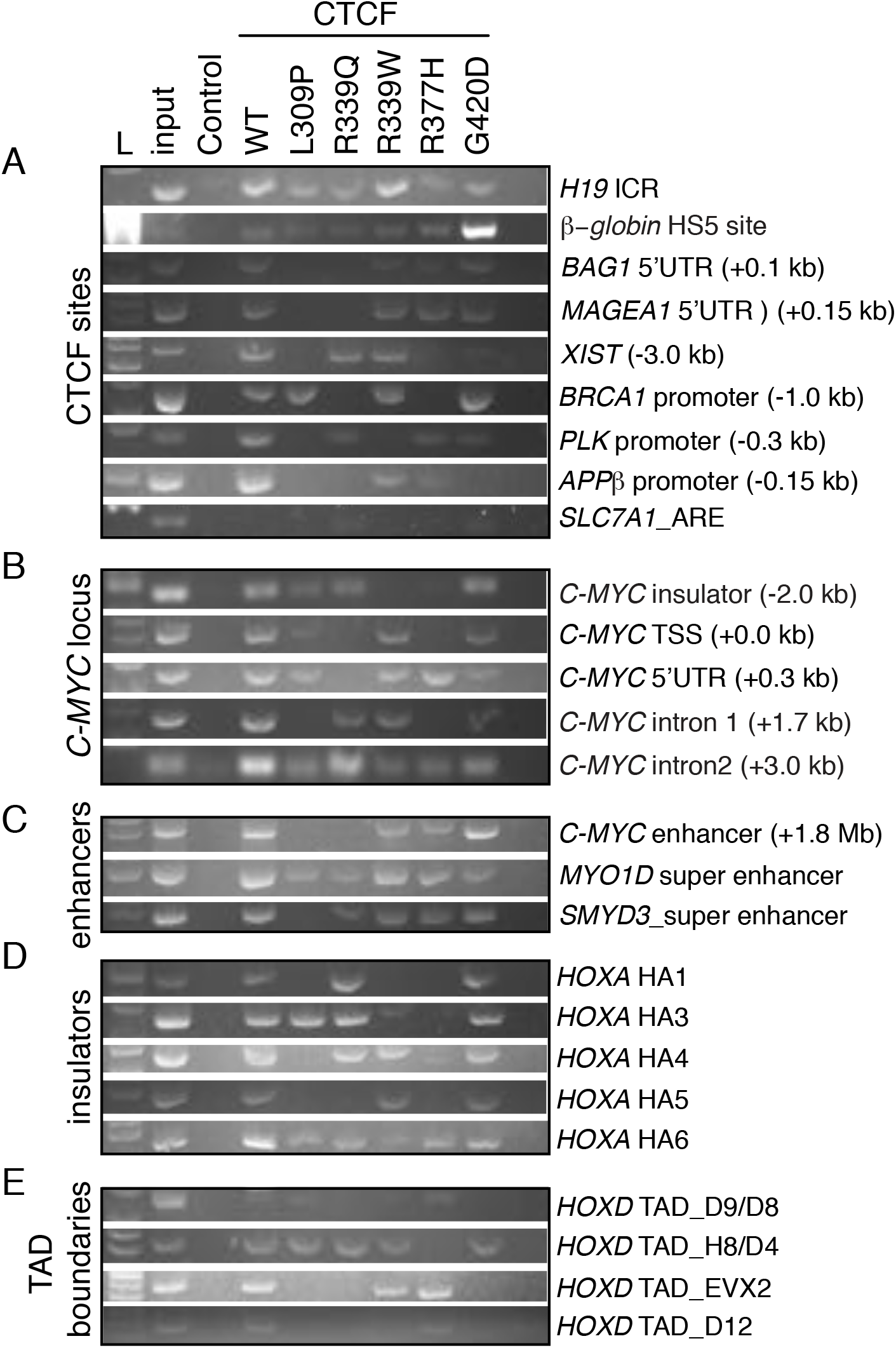
Differential DNA binding exhibited by ZF-mutant CTCF. ChIP-PCR of HA-tagged WT and mutant CTCF expressed in K562 cells; L = 100 bp ladder, input is total genomic DNA before ChIP, Control = eGFP empty vector. Diverse CTCF sites were examined: including **(A)** archetypal CTCF sites; **(B)** the *C-MYC* locus; **(C)** enhancers; **(D)** insulators; and **(E)** TAD boundaries. Where relevant, the genomic distance from the TSS is indicated in brackets. The *SLC7A1* androgen response element (ARE) was used a negative control for CTCF binding. See Supplementary Table 4 listing references for known CTCF sites.

### Molecular dynamics (MD) simulations explain CTCF loss- and gain-of-function ZF mutant phenotypes

To gain insights into the structural impact of these mutations we modelled them on the published crystal structure of CTCF’s ZF domain (ZFs 2-7) in complex with DNA^43^ and performed molecular dynamics (MD) simulations. The locations of the 4 mutated ZF residues were superimposed on the CTCF structure (Figure 5A). The folding free energy change (ΔΔG) calculated for all 5 resulting ZF mutations indicate that each of the mutations are destabilizing. L309P is predicted to have the most severe impact on CTCF folding (ΔΔG = 12.05 kcal/mol), compared to R339Q (ΔΔG = 6.87 kcal/mol), R339W (ΔΔG = 5.00 kcal/mol), R377H (ΔΔG = 5.64 kcal/mol) and G420D (ΔΔG = 1.91 kcal/mol) (Figure 5B). Time evolution studies of secondary structure in WT and mutant CTCF ZF domain indicate that structural elements are stable at the location of each mutation (Supplementary Figure 3). However, β-sheet-forming elements (red) are disrupted by: L309P (ZF2), R339Q, R339W (ZF3) and R377H (ZF4-5) between aa 353 – 363 in ZF4; and R339W, R377H and G420D (ZF6) between 295-305 in ZF2. In all mutants, the β-sheet and turn structure at aa 408 – 418 (ZF6) is also disrupted (Supplementary Figure 3).

**Figure 5.**
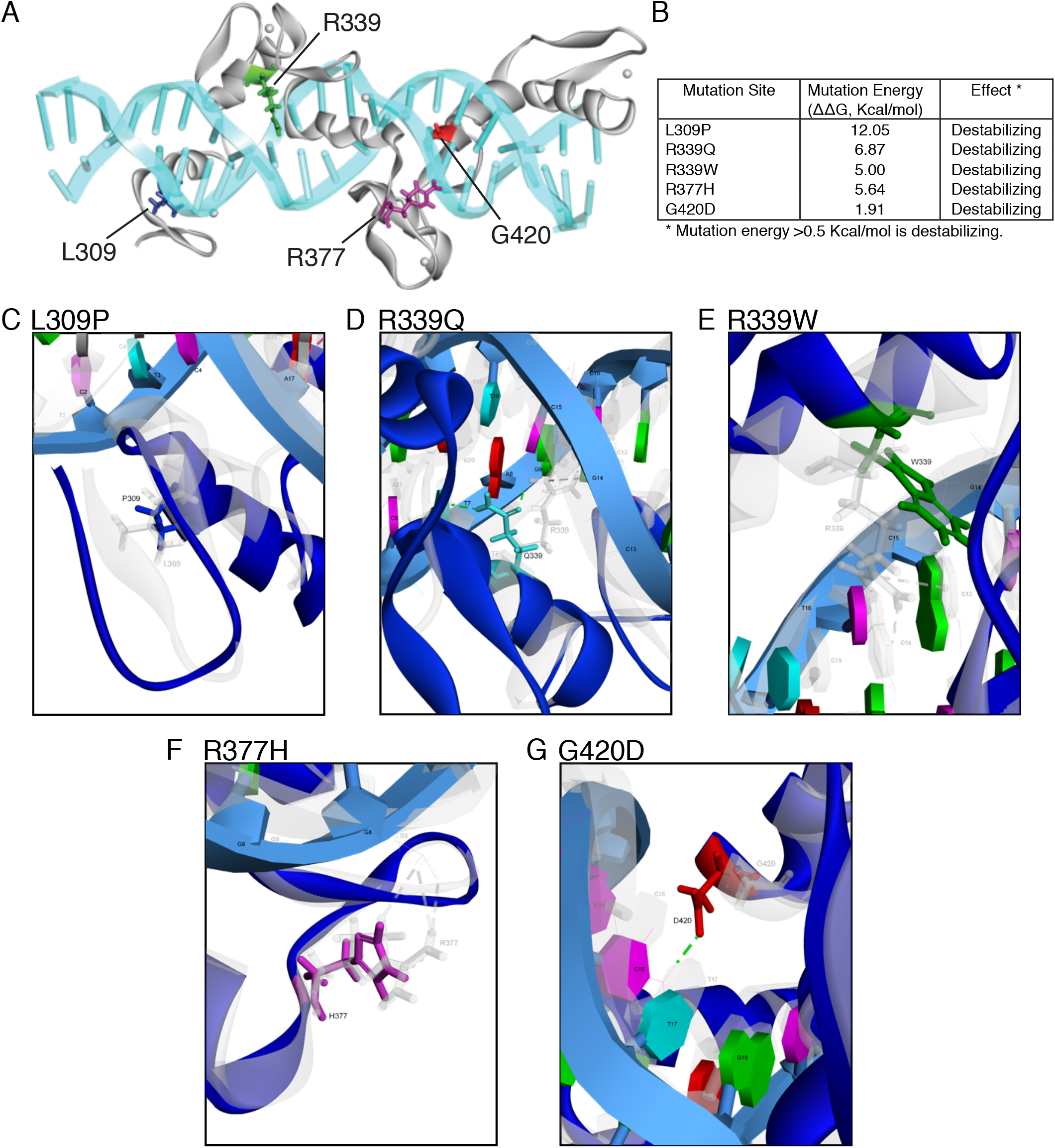
Homology modelling of CTCF ZF mutations. **(A)** CTCF ZF residues impacted by somatic mutation are depicted on the crystal structure model of ZFs 2-7 in association with DNA. Zinc molecules are shown as grey spheres. **(B)** The effect of mutation on the binding energy: the change in minimum free energy (ΔΔG) for WT and mutant CTCF ZF structure in the DNA-bound state. **(C – G)** Overlay images of the normal (WT, grey) and mutant (blue) residues superimposed on the CTCF crystal structure: **(C)** L309P (L grey, P dark blue); **(D)** R339Q (R grey, Q cyan); **(E)** R339W (R grey, W green); **(F)** R377H (R grey, H magenta); **(G)** G420D (G grey, D red). Dashed lines indicate hydrogen bond pairing: old (grey) & new (green). DNA bases and their position relative to the 5’ end of the CTCF consensus are shown.

To examine each mutation in more detail, we examined the superimposed structures of WT and mutant CTCF ZF structures. L309 is facing away from DNA and does not directly contact DNA either before or after mutation to Pro (Figure 5C). Despite this, analysis of molecular interactions between neighbouring CTCF amino acid residues and DNA revealed 7 existing bonds were lost, whereas 12 new bonds were formed (Supplementary Table 3). Root-mean-square deviation (RMSD) measurements showed that the L309P mutation induced a substantial increase in the deviation of the ZF2 backbone compared to WT over the 10 ns simulation run (Figures 6A & B, *p*<0.0001). Similarly, root-mean-square fluctuation (RMSF) measurements spanning the entire ZF 2-7 structure (Supplementary Figure 4) indicated that there was a considerable increase in flexibility (*p*<0.0001, Figure 6C). Consequent to all the conformational changes, the distance of the ZF2 centroid from the DNA centroid was also increased (0.916 Å) in the L309P mutation (Figure 6D).

**Figure 6.**
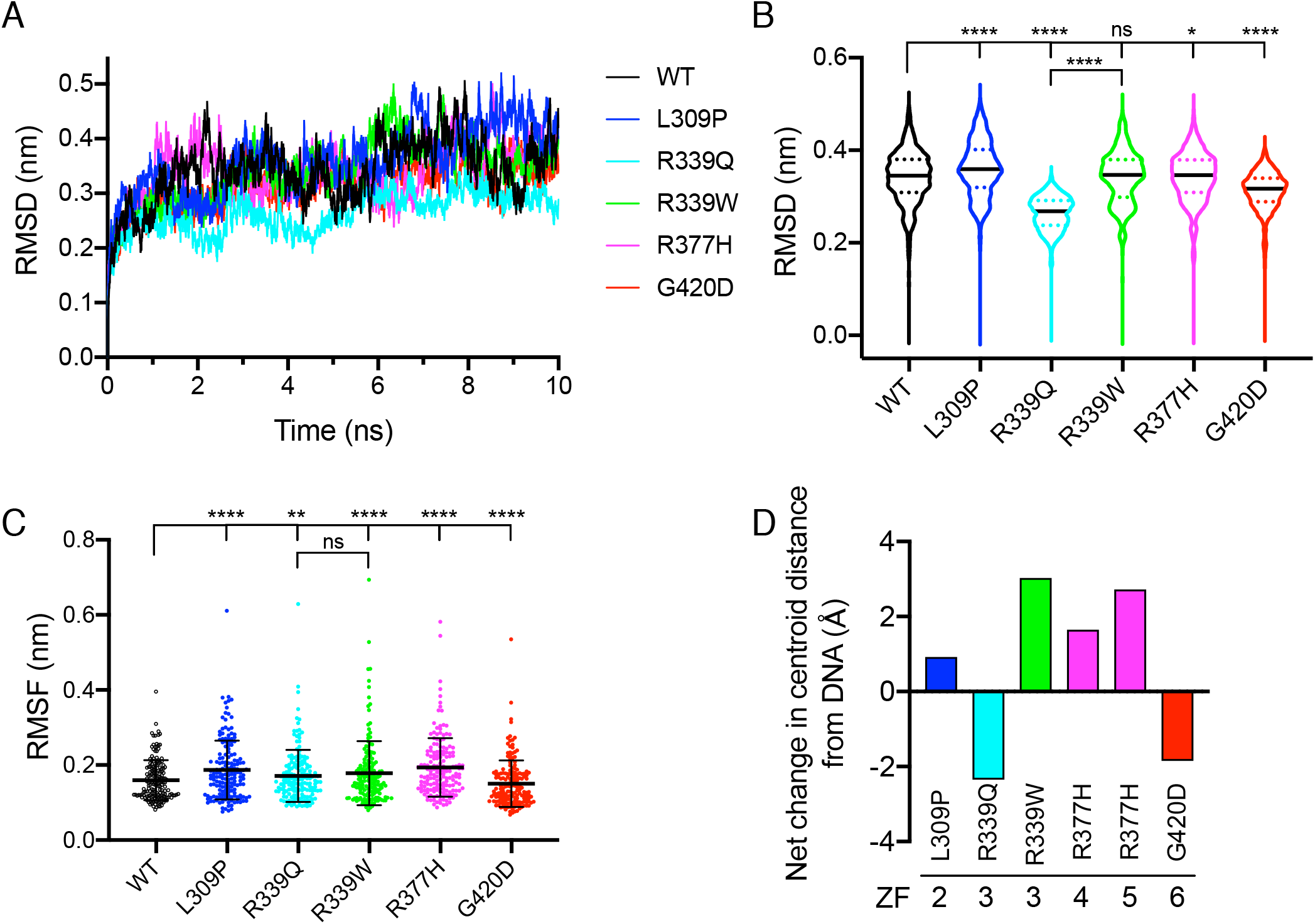
Impact of mutation on CTCF ZF domain conformational stability revealed by MD simulations. **(A-B)** Root-mean-square deviation (RMSD) measurements were calculated from the position differences of backbone atoms in the native (WT) and various mutant conformations. **(A)** Trajectories of all 5 mutants and WT CTCF are displayed over a 10 ns time span, measured at 2 ps intervals. **(B)** Violin plots of all RMSD measurements (0.000 −10.000 ns, 5001 in total). In each plot, the solid black line indicates median and dashed coloured lines indicate quartiles. **(C)** Root-mean-square fluctuation (RMSF) measurements were obtained for all residues (n=173) at each time point for the WT and mutant structures. In each plot, the mean ± SD is shown. In **(B)** & **(C)** the Wilcoxon matched-pairs signed rank test was applied to all paired measurements (ns = not significant; *, *p*<0.05; **, *p*<0.01; ****, *p*<0.0001). **(D)** The net change in distance (in Å) of the centre of mass (centroid) of the associated ZF domain from DNA compared to WT CTCF.

Arginine 339 at DNA-binding position ‘+6’ within ZF3 directly contacted guanine (G14) and cytosine (C13) residues on one DNA strand via two hydrogen bonds and one cation-π bond, however mutation to Q (R339Q) or W (R339W) abolished these bonds (Figure 5D & E, Supplementary Table 3). Remarkably, Q339 formed two new hydrogen bonds: firstly, between the Gln side chain carbonyl group and cytosine (C15); and secondly, between the side chain amide group and thymine (T7) on the complementary strand (Figure 5D). Both mutations also disrupted the interaction of E336 with cytosine (C15), with 6 and 4 new bonds formed at neighbouring residues for Q339 and W339 respectively (Supplementary Table 3). MD simulations showed that the R339Q triggers less conformational deviation than WT or R339W (Figures 6A & B, both *p*<0.0001), however over the entire ZF structure R339Q and R339W both exhibited more flexibility than WT (Figure 6C, *p* = 0.0018 & *p*<0.0001 respectively). Consequently, R339Q shifted ZF3 towards DNA (2.342 Å) and in the case of R339W, ZF3 moved away from DNA (3.021 Å) (Figure 6D).

R377H, which occurs in an inter-ZF residue between ZF4 and ZF5, disrupted three hydrogen bonds that stabilise the interaction of R377 with the DNA phosphate moiety at guanine (G8) (Figure 5F). Adding to this, 22 neighbouring molecular contacts are lost and 22 new bonds are formed (Supplementary Table 3). RMSD measurements show that R377H induced an increased deviation in the conformation over time (Figures 6A & B, *p* = 0.0439) and increased flexibility in the entire ZF structure (Figure 6C, *p*<0.0001). As a result, both ZF4 and ZF5 were shifted away from the DNA phosphate backbone (1.643 Å & 2.718 Å respectively, Figure 6D).

Finally, CTCF modelling confirms glycine at position 420 and DNA-binding position ‘+2’ in ZF6 does not directly contact DNA (Figure 5G). However, when mutated to aspartic acid (G420D), a new hydrogen bond is formed between the side chain carbonyl group and cytosine (C16) in the core consensus sequence (Figure 5G). A net loss of 4 bonds at neighbouring residues was also observed (Supplementary Table 3). RMSD measurements showed that G420D exhibited decreased structural deviation during the simulation run (Figures 6A & B, *p*<0.0001) and decreased RMSF values compared to WT indicating reduced flexibility (*p*<0.0001, Figure 6C). Consequently, G420D resulted in ZF6 shifting 1.841 Å toward the DNA (Figures 6D).

In summary, our data suggest that mutations R339W and R377H disrupted CTCF’s primary interactions with DNA and, along with the highly destabilising L309P mutation, are responsible for shifting ZF domains away from DNA. Importantly, R339Q and G420D both formed new primary bonds and the associated ZF domain moved nearer to the DNA.

## Discussion

Tumour-specific mutations in CTCF were described almost two decades ago^34^. In that seminal report, functional testing of some ZF mutations comprised electrophoretic mobility shift assays (EMSA) and a reporter assay. The key finding from these preliminary functional studies was that these somatic CTCF ZF mutations exhibited selective disruption in DNA binding to some, but not all, CTCF target sites, giving rise to the concept of ‘change-of-function’ mutations^34^. Since then the functional characterisation of CTCF ZF mutations has been limited, despite many landmark cancer genome studies reporting hundreds of missense somatic CTCF ZF mutations.

The potential genome-wide impacts of CTCF ZF mutation on DNA binding were examined through mutation in all 11 ZFs of a key Zn^2+^ co-ordinating histidine residue within the conserved C2H2 tetrahedron arrangement that co-ordinates ion binding^15^. This approach, whilst not directly emulating tumor-specific missense mutations, revealed that in all cases DNA binding was not completely abolished. Indeed, residual DNA binding ranged from ~15-80% depending on the position of the ZF^15^. Our previous report showed that the 3 CTCF ZF mutations most frequently occurring in endometrial cancers (K365T, R377H, P378L) had differing impacts on CTCF function when overexpressed^38^. The two inter-ZF mutations (R377H and P378L) abrogated CTCF’s anti-proliferative and anti-clonogenic effect in Ishikawa endometrial cancer cells. However, mutation of K365, a key DNA-binding residue at position ‘+3’ in ZF4, to threonine, had no impact on proliferation or colony formation^38^, despite causing a 20-fold loss of CTCF DNA-binding affinity^43^. Interestingly, CTCF K365T conferred significantly increased resistance to UV-induced apoptosis in Ishikawa cells compared to WT CTCF, suggesting the first pro-tumourigenic role of CTCF ZF mutations^43^.

Initial deletion mutagenesis studies of the CTCF ZF domain indicated ZFs 3-11 were required to bind the human *c-myc* promoter^46^. Further refinement was achieved in two studies, in which similar binding modes were confirmed on the chicken *c-myc* promoter requiring central ZFs 2-7^46^ or ZFs 3-8^47^. Furthermore, the central ZFs 5-7 of CTCF were required for *APPβ* promoter binding, however it was the peripheral ZFs which provided the stability in DNA binding^47^. Whilst gel mobility shift analysis confirmed that CTCF ZFs 4-7 were necessary and sufficient to bind to a core 12 bp consensus sequence^48^, the first crystal structure of the CTCF ZF domain resolved that ZFs 3-7 bound the 15 bp core DNA consensus sequence^43^. Nakahashi *et al* proposed a ‘saddle’ model containing a core CTCF motif (C) bound by central ZFs 4-7 as well as upstream (U) and downstream (D) modules bound by peripheral ZFs^15^. Yin *et al* further refined this model by describing 4 CTCF binding site modules^49^. Modules 4, 3 and 2, spanning the 15 bp core CTCF consensus as well as downstream sequences, are bound by ZF3, ZFs 4-6 and ZF7 respectively; whilst upstream module 1 is bound by ZFs 9-11^49^. These studies provide insights as to why mutations occurring in different CTCF ZFs may produce diverse effects on DNA binding and functional outcomes depending on where in the modular binding mode the mutant ZF residue occurs.

The five different somatic missense ZF mutations we examined in this study each occur in key positions within the central zinc fingers. Each mutation provided critical insights into CTCF structure-function relationships. The spatial arrangement of residues within the C2H2 ZF finger motif, including the flexible inter-ZF link, are critical to maintaining ZF structure, and are therefore very highly conserved^50^. Somatic ZF mutations did not affect overall CTCF protein stability or localisation within the nucleus when stably expressed. However, when transiently overexpressed, CTCF ZF mutants clearly decreased transcriptional activation compared to WT CTCF. Interestingly, our previous endometrial cancer study indicated that missense ZF-containing *CTCF* alleles were expressed at a higher frequency than WT alleles, when comparing RNA sequencing to DNA sequencing^38^. This suggested that expression of aberrant CTCF was up-regulated to functionally compensate for a deficit in CTCF function. In our study, we observed differing impacts on CTCF-mediated cellular proliferation. These impacts on CTCF function are attributable to a change or gain in DNA-binding specificity.

L309 typifies an invariant hydrophobic residue (always Leu or Met) in the alpha helical region of all CTCF C2H2 ZF fingers. Residues near the C-terminal end of each C2H2 ZF fold into an alpha helix, positioning key amino acids within the helix to interact directly with DNA^50^. L309 mutation to Pro (L309P) affected the thermodynamic stability of ZF2, most likely through the α-helical-breaking tendency of proline in water-soluble proteins. We confirmed increased RMSD and RMSF values during molecular dynamics simulations and a shift of CTCF away from DNA. Not surprisingly, despite some DNA-binding still being maintained, L309P exhibited loss-of-function in *in vitro* cell growth assays. R339 mutation to Q or W differentially impacted CTCF growth and colony forming function and this too was explained by molecular dynamics simulations. For R339Q, the ZF domain of the CTCF shifts closer to DNA and two new hydrogen bonds are formed at Q339, explaining the intermediate loss-of-function cell growth phenotype observed with R339Q. R339W, which exhibits loss-of-function phenotypes, disrupts all primary DNA contacts and deflects ZF3 away from DNA. This is despite R339W still maintaining nearly half (47%) of DNA binding genome-wide^15^.

R377 resides in one of the inter-ZF regions which act as bridges between ZFs to allow flexibility in the DNA-free form but stability in the DNA-bound form^43^. Our modelling showing that R377 contacts the DNA phosphate backbone was also confirmed by structural studies of the CTCF ZF domain bound to the *Pcdh* enhancer^49^. Hence, not all amino acids at canonical DNA binding positions in ZFs directly contact DNA, such that intra- and inter-ZF residues are also involved in DNA contacts^49^. The R377H mutation, which eliminates this DNA interaction, also destabilises neighbouring molecular interactions. R377H exhibited loss-of-function cell growth phenotypes in K562 erythroleukaemia cells, similar to our observations in endometrial cancer cells^38^. Remarkably, despite G420D exhibiting some loss of binding to target sites and loss of gene regulatory activity, a gain-of-function was observed as it suppressed colony formation to a greater extent than WT CTCF. Consistent with these phenotypes, G420D formed a new bond with DNA and resulted in ZF6 shifting towards DNA. Overall, these studies reveal that the location of the ZF missense mutation determines the impact it has on loss-, change- or gain-of-function. We examined mutations in DNA-contacting residues, in a residue co-ordinating ZF folding, in the inter-ZF region and within or outside the central core consensus binding ZFs 4-7. Furthermore, the mutant amino acid residue side chain can also have a significant impact on cellular phenotypes.

Different ZF modules with identical DNA specificity residues (at positions −1, +2, +3 and +6) can bind different sequences, influenced by DNA sequence context and inter-ZF residue-residue interactions^51^. Furthermore, neighbouring ZFs may affect the DNA-binding conformation and specificity of a particular ZF^52^. Our data has revealed that those residues in close proximity to the mutant residue can lose existing bonds or acquire new DNA interactions. Missense ZF mutations in CTCF can destabilise the DNA-bound conformation of neighbouring ZFs. Thus, MD simulations have illuminated the broad and diverse impact that CTCF ZF mutations exert on DNA binding.

Human C2H2 ZF-containing TFs contain on average 10 C2H2-ZF domains, leading to possible binding sites of up to ~30 bp. These suffice to specify target sites in the human genome^14^. However, not all CTCF ZFs bind DNA simultaneously, meaning some may be oriented for RNA or protein-protein interactions. Thus, the impact of somatic CTCF ZF missense mutations on: i) RNA interactions; ii) protein-protein interactions; and iii) higher-order chromatin structure has yet to be examined. CTCF is also a high affinity RNA-binding protein conferring long-range chromatin interactions mediated via RNA^53^. *Jpx*, a long non-coding RNA known to activate *Xist*, interacts with CTCF and relieves CTCF-mediated repression at the *Xist* locus, suggesting that *Jpx* directly interferes with CTCF DNA-binding via its ZF domain^54^. In CTCF, ZFs 10 and 11 in concert with the C-terminus have been defined as an RNA-binding region, binding the long noncoding RNA *Wrap53*^55^. Mutations in peripheral ZFs 1 and 10 can disrupt CTCF interactions with RNA, leading to disruption of CTCF-mediated chromatin looping and higher-order chromatin structure^55^.

Most C2H2-containing proteins interact with a diverse range of proteins, defining their role as transcriptional activators or repressors^13^. This confers additional gene regulatory complexity at specific DNA sites, potentiating the disruption of locus-specific gene expression as a result of ZF mutations. CTCF homodimerisation, which is essential for correctly orientating CTCF DNA looping and topology, is orchestrated via inter-CTCF interactions between the ZF domain and C-terminus^56^. The CTCF ZF domain interacts with TFs YB1, OCT and SIN3A as well as the chromatin remodeler CHD8^57^, amongst other proteins. The proteome-wide impact of somatic CTCF ZF missense mutations on homodimerisation and protein-protein interactions is yet to be elucidated.

Finally, it is not known what impact somatic ZF missense mutations will have on rewiring genomic architecture. Topologically associated domains (TADs) are discrete territories, compartmentalising the genome into independent, often evolutionarily conserved domains^20,21,23,58,59^. TADs are characterized by frequent CTCF-mediated contacts within domains and a low frequency of contacts between domains^60^. These TADs are themselves demarcated into sub-megabase sub-TADs and loop domains, often differentially co-ordinated by CTCF interaction with other architectural proteins such as cohesin^19,24,61^. Deletion or inversion of CTCF sites at TAD boundaries can drastically affect gene regulation, leading to ectopic activation of gene expression due to illegitimate promoter and enhancer interactions, often with pathogenic consequences^26,27,62,63^. In cancer, genetic alteration or hypermethylation of CTCF sites at TAD boundaries can disrupt chromatin topology and lead to aberrant activation of oncogenes^64–66^. The global impact of somatic missense mutations in CTCF, which typically only occur on one allele and cause *CTCF* haploinsufficiency, remains to be investigated.

## Conclusions

Over the last decade, unprecedented insights into CTCF’s essential role in genome organisation and architecture have been revealed via the generation of high-resolution maps of chromatin interactions by chromosome conformation capture (3C)-based techniques. However, the structure-function relationships of CTCF mutations, particularly within the DNA-binding ZF domain, have not been investigated. We reveal that the CTCF ZF domain is significantly mutated in cancer, with ZF-specific missense mutations impacting CTCF’s anti-proliferative capacity, DNA-binding and gene regulatory activities. Strikingly, we observed a broad spectrum of functional impacts ranging from complete, partial or no loss-of-function in cellular growth phenotypes and transcriptional regulation, as well as gain-of-function, resulting from the formation of new bonds between the mutant ZF and DNA. Our MD simulations revealed that all CTCF ZF mutations were destabilising, with the loss or gain in DNA binding not just localised to the mutant residue. This highlights the importance of understanding structure-function relationships in normal and mutated CTCF. As *CTCF* exhibits haploinsufficiency in cancer, the interplay between mutant and wildtype CTCF at specific loci and at target sites genome-wide remains an unanswered question. Understanding how somatic CTCF ZF mutations affect RNA, protein-protein or chromatin interactions globally will be the next frontier in understanding the molecular pathophysiology of cancer.

## Materials and Methods

### Cell lines

Human erythroid leukaemia (K562) cells were grown in RPM1 1640 medium and human embryonic kidney (HEK293T) cells were cultured in DMEM. All basal media were supplemented with 10% FCS (v/v), penicillin (100 U/mL) and streptomycin (100 μg/mL). All human cell lines were previously authenticated by short tandem repeat profiling (Cellbank, Australia).

### Reagents: Expression Vectors and Antibodies

The lentiviral vector pCCLteteGFP2AHAhCTCF^33^ was used to express CTCF ZF mutations. PCR amplicons containing ZF mutations (L309P, R377H, G420D) were generated by splice overlap extension PCR and were cloned in using *BmgBI/ClaI*. R339Q and R339W mutations were created by gene synthesis from DNA2.0 and sub-cloned using *Bsu36I/Tth111I*. Y226C and R603C were synthesized as Geneblocks (IDT) and cloned into *BstX1*/*BstXI* sites and *PstI/BlpI* sites respectively. Primary antibodies include: CTCF rabbit monoclonal (#3418, Cell Signaling Technology; 1:5,000), HA epitope mouse monoclonal (HA.11, Covance; 1:5,000) and GAPDH (ab8245, Abcam; 1:5,000). Secondary antibodies include: rabbit or mouse antibodies conjugated to horseradish peroxidase (HRP, Millipore; 1:5,000).

### Retroviral and lentiviral transduction

Viral supernatants were produced by calcium phosphate transfection of HEK293T cells: with pJK3, pCMVTat and pL-VSV-G packaging plasmids used to produce replication-defective retroviruses; and pRSV-Rev, pMDLg/p.rre and pMD2.VSV-G used to produce replication-defective lentiviruses. Viral supernatants collected after 24-48 h were 0.45 μM-filtered and snap-frozen or concentrated by ultracentrifugation for 2 h at 20,000 rpm in a SW28 Beckman rotor. Viral supernatant was resuspended on ice in 10% (v/v) FCS/DMEM at 1/100th of the original volume. Attached cells (1-5×10^5^/well) were seeded in 6-well plates before addition of fresh medium containing viral supernatant (~5×10^5^ transducing units) and Polybrene (8 μg/mL; Sigma) and ‘spin-oculated’ for 90 min at 1,500 rpm. The supernatant was replaced with medium 12 h post-transduction and fluorescent cells were purified 24 h later by fluorescence-activated cell sorting (FACS; >95% purity on re-analysis) using a FACS Influx (Becton Dickinson, BD). K562 cells (~5×10^5^/mL) in 1 mL medium with 4 μg/mL Polybrene were placed in a 5 mL capped FACS tube and transduced with viral supernatant for 90 min by ‘spin-oculation’. The cells were resuspended, incubated at 37°C for 4 h before removal of viral supernatant. For *in vitro* assays, cells were either plated out immediately or allowed to recover after sorting for 48 – 72 h in medium containing 100 μg/mL Normocin (Invivogen).

### Immunofluorescence

Transduced K562 cells (1 × 10^6^) were fixed with equal volume of 4% (w/v) formaldehyde for 20 min at room temperature (RT). Cells were centrifuged at 2,000 rpm for 3 min and resuspended in PBS twice. Cells (5.0 × 10^5^) were allowed to settle onto coverslips coated with poly-D-lysine, before drying and permeabilisation with Triton X-100 0.5% (v/v) in PBS for 10 min at RT. Cells were rinsed three times in PBS and blocked in 3% (w/v) BSA/PBS for 40 min at RT. Cells were rinsed three times in PBS and incubated with mouse anti-HA antibody (1:500, HA.11, Covance) for 90 min at RT. Cells were rinsed three times in PBS and incubated with F(Ab’)2-goat anti-mouse IgG-Alexa 594 (1:500, #A11020, ThermoFisher Scientific) and DAPI (1:10,000, #D1306, Life Technologies) for 40 min at RT. Cells were rinsed three times in PBS and mounted using Prolong Gold Antifade (Life Technologies). Slides were imaged at 60x using the DeltaVision Personal (Applied Precision) and the DAPI, FITC and A594 filters. Images were analysed after deconvolution using Volocity software.

### Western blot analysis

Protein extracts were prepared with cell lysis buffer containing 20 mM TrisCl (pH 8), NaCl (150 mM), 1% (v/v) Triton X-100, 0.1% (v/v) SDS, 0.5% (w/v) sodium deoxycholate and EDTA-free protease inhibitor cocktail (cOmplete, Roche Life Science), prior to separation using denaturing sodium dodecyl sulfate polyacrylamide gel electrophoresis (SDS-PAGE). Proteins were transferred in a semi-dry transfer apparatus to PVDF membrane before immunoblotting. Membranes were blocked in PBST containing 20% (v/v) BlokHen (AvesLab) or PBST containing 0.3% (w/v) BSA, 1% (w/v) polyvinylpyrrolidone, 1% PEG (mw 3350). Protein expression was detected using primary antibodies followed by washing and staining with appropriate secondary antibodies against rabbit, goat or mouse IgG conjugated to horseradish peroxidase (HRP). The HRP substrate SuperSignal^®^ Chemiluminescent Substrate (Pierce) was detected on a Kodak Imagestation 4000R Pro or BioRad Chemidoc Touch. Blots were stripped with ReBlot Plus (Merck Millipore) prior to re-probing with protein loading control antibodies.

### Mutation and Bioinformatic analysis

CTCF mutations were obtained from the Catalogue of Somatic Mutations in Cancer (COSMIC) portal, The Cancer Genome Atlas (TCGA) cBIO portal and published reports (see Supplementary Table 1). Single nucleotide variants were obtained from dbSNP. The potential impact of mutations was determined using Polyphen-2. All amino acid alignments were performed using the Clustalw algorithm within MacVector. A raw alignment of CTCF ZFs was exported into Weblogo (weblogo.berkeley.edu/logo.cgi) to generate a sequence logo. The maximum sequence conservation for an amino acid is log_2_20 ~ 4.32 bits. CTCF target sites in CTCF-expressing plasmid pCCLteteGFP2AHA-hCTCF were predicted using MatInspector (Genomatix).

### Cell biological assays

Cell proliferation was assessed by 3-(4,5-^1,2^methylthiazol-2-yl)-2,5-diphenyltetrazolium bromide (MTT) assay (Merck Millipore). K562 cells (5,000/well) were plated in triplicate in a 96-well plate and proliferation was assessed over 4 d by the addition of MTT at 37°C overnight. The reaction was quenched with isopropanol/HCl and then absorbance was measured at 572 nm using a Wallac 1420 Victor plate reader (Perkin Elmer). The clonogenic capacity of K562 cells was measured by plating 5,000 cells diluted in Iscove’s Modified Dulbecco Medium (Life Technologies) containing 3 mL Methocult GF H4230 (Stem Cell Technologies) and plated in triplicate in 35 mm gridded plates and incubating for 8-10 d.

### Chromatin Immunoprecipitation

K562 cells (1×10^6^ in 1 mL) were transduced with 10-60 uL viral supernatant of the control (eGFP only), CTCF WT and five CTCF mutants. After 72 h the cells were assessed by flow cytometry (LSR Fortessa, Becton Dickinson) and shown to vary between ~14-21% expression. For each chromatin immunoprecipitation (ChIP), 5×10^6^ cells were cross-linked with 1% (w/v) formaldehyde for 10 min and then quenched with 1 M glycine to a final concentration of 20 mM. Nuclear lysates were sonicated for 25 cycles, 30 s on, 30 s off using a Bioruptor sonicator (Diagenode). For each immunoprecipitation, 3 μg of rabbit polyclonal antibody against the HA epitope (ab9110, Abcam) was used. Magna ChIP ™ Protein A/G conjugated magnetic beads (Millipore) were used to immunoprecipitate antibody-bound chromatin complexes, and all subsequent steps were performed according to the manufacturers’ instructions. After de-crosslinking, phenol/chloroform extraction, and ethanol precipitation, PCR was performed on genomic DNA targets using Phusion polymerase with GC buffer (Finnzyme). PCR primers spanning experimentally validated CTCF targets sites were designed from previous reports (see Supplementary Table 4). A more detailed protocol is available on request.

### Transfection of WT and mutant CTCF

HEK293T cells (1×10^5^) were plated into 12-well plates, 18 h before transfection. In each transfection, pCCLteteGFP2AHACTCF WT, mutant or empty vector (0.5 μg) was combined with 2 μL Lipofectamine 2000 (ThermoFisher) in OptiMEM medium (Gibco) according to the manufacturers’ instructions. After 48 h, cells were detached and assessed for eGFP by flow cytometer (LSR Fortessa, Becton Dickinson) and then lysed with cell lysis buffer.

### Structural Modelling and MD simulations

A 3.2 Å X-ray diffraction crystal structure representing the CTCF ZFs 2-7 / DNA complex (PDB: 5T0U)^43^ was used as the initial template to prepare CTCF mutant models. The template was optimized using ‘Prepare Protein’ and ‘Energy Minimization’ protocols available in Biovia Discovery Studio (DS) 2017R2 software suite. Initial mutant models (L309P, R339Q, R339W, R377H and G420D) were built using ‘Build and Edit Protein’ tool in DS by substituting original amino acid residues for the respective mutant. These mutant models were further optimized for their minimum energy confirmation using steepest descent algorithm in DS with a non-bonded lower cut-off distance of 10 Å. Impact of mutations on protein stability was analyzed using ‘Calculate Mutation Energy (Stability)’ protocol in DS. The protocol calculates the difference in the free energy of folding (ΔΔG_mut_) between the mutant structure and the wild type protein as follows:

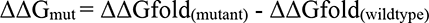

- where ΔΔGfold is defined as the free energy difference between the folded and unfolded state of the protein. The unfolded state is modelled as a relaxed protein in extended conformation with the mutated residue in the center.

To further analyse the impact of the mutation on CTCF binding ability to DNA, we first superimposed mutant CTCF models onto the wildtype CTCF model in complex with DNA using Chimera version 1.14. Furthermore, we performed MD simulations of WT and all the mutant models using GROMACS version 4.5.3. The system (CTCF mutant model in complex with DNA) was placed in the center of a cubic box with at least 1 nm from the box edges. The ions (Na^+^ and Cl^−^) were added to the system for neutralizing and preserving at physiological concentration (0.15 M). The protocol consisted of successive rounds of energy minimization, annealing, equilibration, and trajectory production in implicit solvent represented by the generalized Born/solvent-accessible surface area model and using a distance cut-off of 10 Å to short-range, non-bonded interactions. Keeping backbone atoms restrained, the protein was relaxed with 50,000 steps of energy minimization, followed by annealing with a 60 – 300 K temperature ramp applied over 100 ps. In equilibration, the temperature was maintained at 300 K using Langevin dynamics for 50,000 steps over 100 ps. Production simulations were performed in the isothermal-isobaric ensemble, keeping both the DNA fragment and CTCF unconstrained. Bonds between hydrogen and heavy atoms were constrained at their equilibrium length using LINCS algorithm. Root-mean-square deviation (RMSD), root-mean-square fluctuation (RMSF), secondary structure and interaction energy analyses were carried out using GROMACS for the entire simulation run. All non-bonded interactions for the final poses of CTCF wildtype and mutants in complex with DNA were identified using DS.

## List of Abbreviations

ChIP: chromatin immunoprecipitation
CTCF: CCCTC-binding factor
DMEM: Dulbecco’s Modified Eagle Medium
EMSA: electrophoretic mobility shift assay
FACS: fluorescence-activated cell sorting
HA: hemagglutinin
ICR: imprinting control region
MD: molecular dynamics
MTT: 3- (4,5-^1,2^methylthiazol-2-yl)-2,5-diphenyltetrazolium bromide
PBST: phosphate buffered saline Tween 20
PDB: Protein Data Bank
RMSD: root-mean-square deviation
RMSF: root-mean-square fluctuation
RPMI: Roswell Park Memorial Institute
SNP: single nucleotide polymorphism
TAD: topologically associating domains
TF: transcription factor
WT: wildtype
ZF: zinc finger.

## Declarations

### Ethics approval and consent to participate

Not applicable.

### Consent of publication

Not applicable.

### Availability of data and material

All data generated or analysed during this study are included in this published article (and its supplementary information files). Additionally, PDB files for CTCF ZF WT and mutant conformations have been submitted to Protein Data Bank, accession files: L309P, 1A2B; R339Q, 3C4D; R339W, 5E6F; R377H, 7G8H; G420D, 9I0J.

### Authors’ contributions

C.G.B conceived the study, designed constructs, analysed data and wrote the manuscript. J.E.J.R. reviewed the manuscript, supervised research governance and provided scientific discussion; C.M., P.O’Y., M.A., T.L. and H.F. performed cellular biology experiments; W.K. and T.L. performed ChIP and preliminary structural modelling; C.S. performed immunofluorescence; C.G.M provided mutation data; U.S. analysed mutation data and reviewed the manuscript; S.G & K.P.S & O.W. performed and reviewed structural modelling, docking and molecular dynamics simulations.

## Acknowledgements

The authors acknowledge The University of Sydney High Performance Computing service at The University of Sydney for providing resources that have contributed to the research data reported within this paper.

## Authors’ information

Not applicable

## Supplementary Figure Legends

**Supplementary Figure 1.**
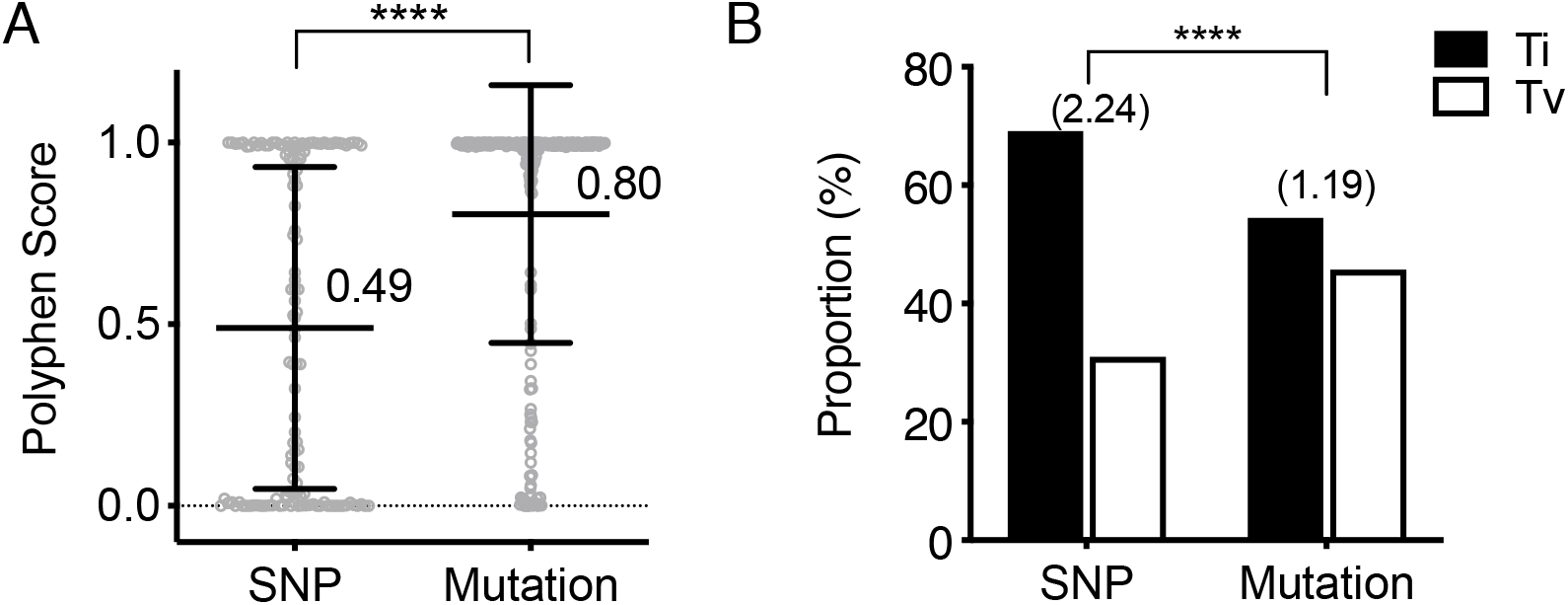
Predicted phenotypes of missense CTCF SNPs and somatic mutations. **(A)** The mean Polyphen score of missense SNPs and somatic mutations in CTCF is shown. **(B)** Proportion of the transition and transversion mutations for SNPs and somatic missense mutations in CTCF; ratio in brackets. Data represent the mean ± SD with statistical analysis performed using Mann-Whitney U-test **(A)** or Chi-square test **(B)** (****, *p*<0.0001).

**Supplementary Figure 2.**
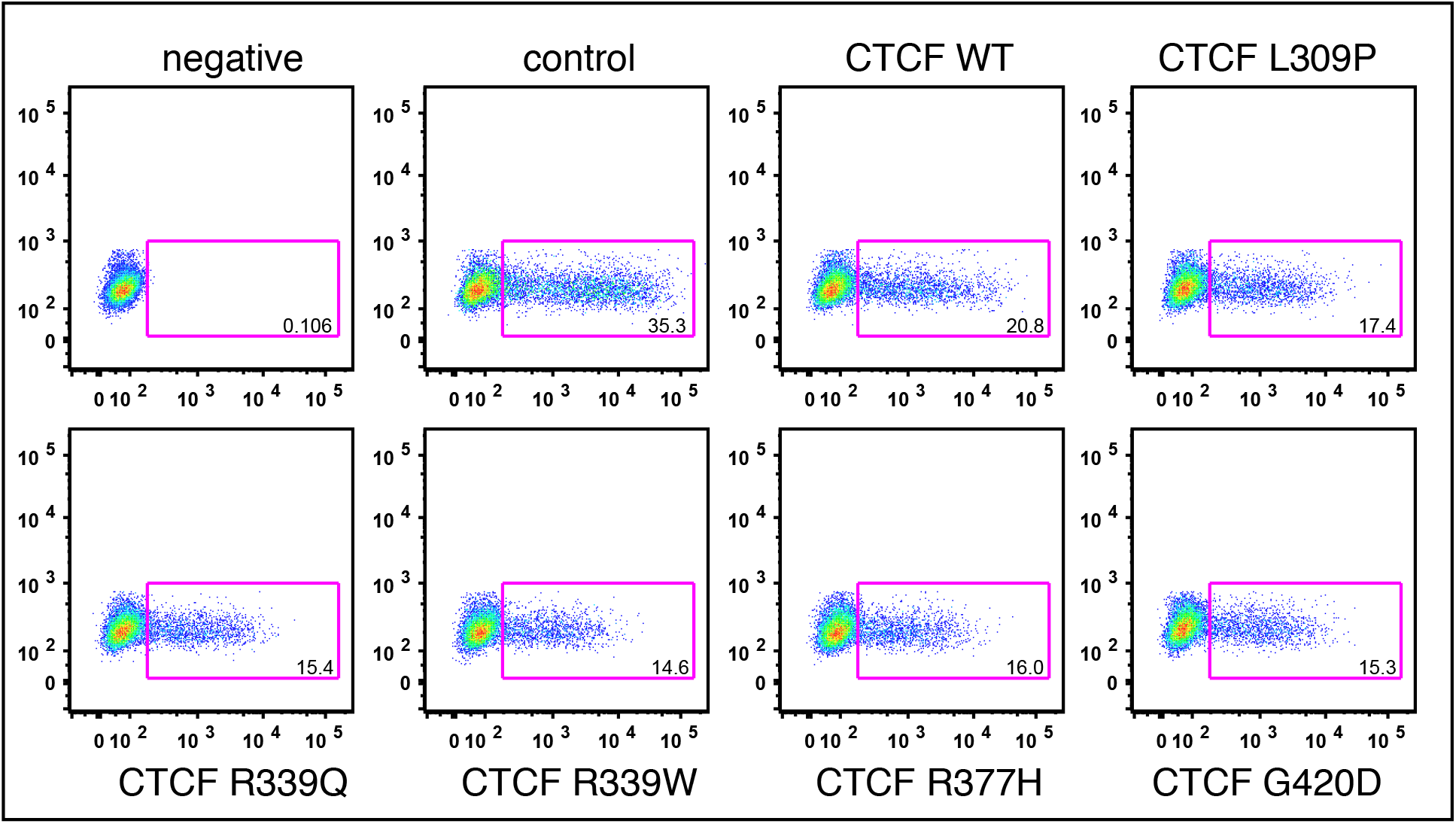
WT and mutant CTCF expression in K562 cells. Flow cytometry of eGFP expression achieved from transduction of HA-tagged WT and mutant CTCF lentiviral vectors in K562 cells. Cells were lysed for immunoblot (Figure 2B), prepared for immunofluorescence (Figure 2C) and subjected to formaldehyde cross-linking for ChIP (Figure 4).

**Supplementary Figure 3.**
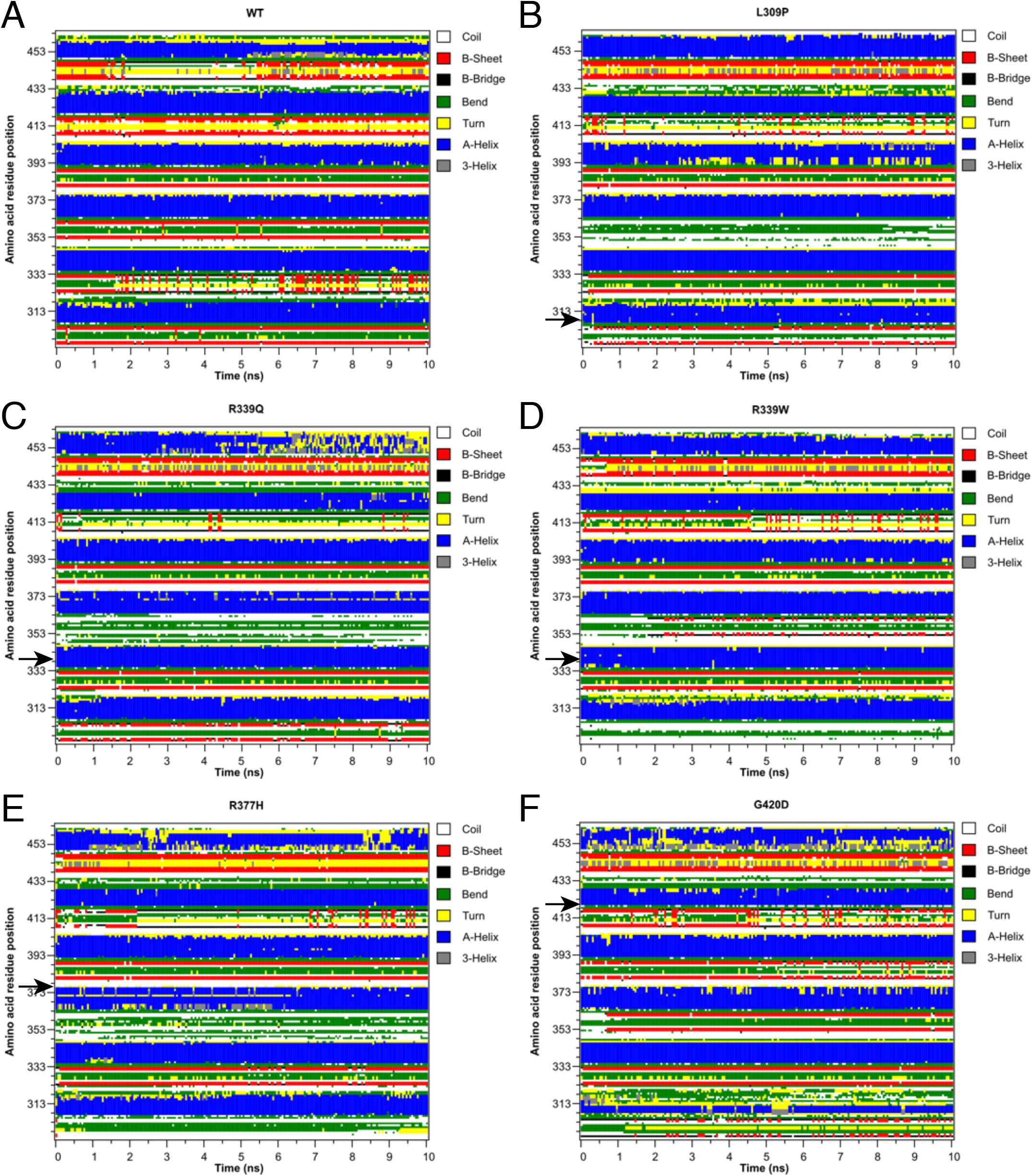
Time evolution of secondary structure in WT and mutant CTCF ZF structures. The Dictionary of Secondary Structure of Proteins (DSSP) classification of secondary structure was calculated for all amino acid residues over the 10 ns time course: **(A)** WT; **(B)** L309P; **(C)** R339Q; **(D)** R339W; **(E)** R377H; and **(F)** G420D. Arrows indicate the position of each mutation.

**Supplementary Figure 4.**
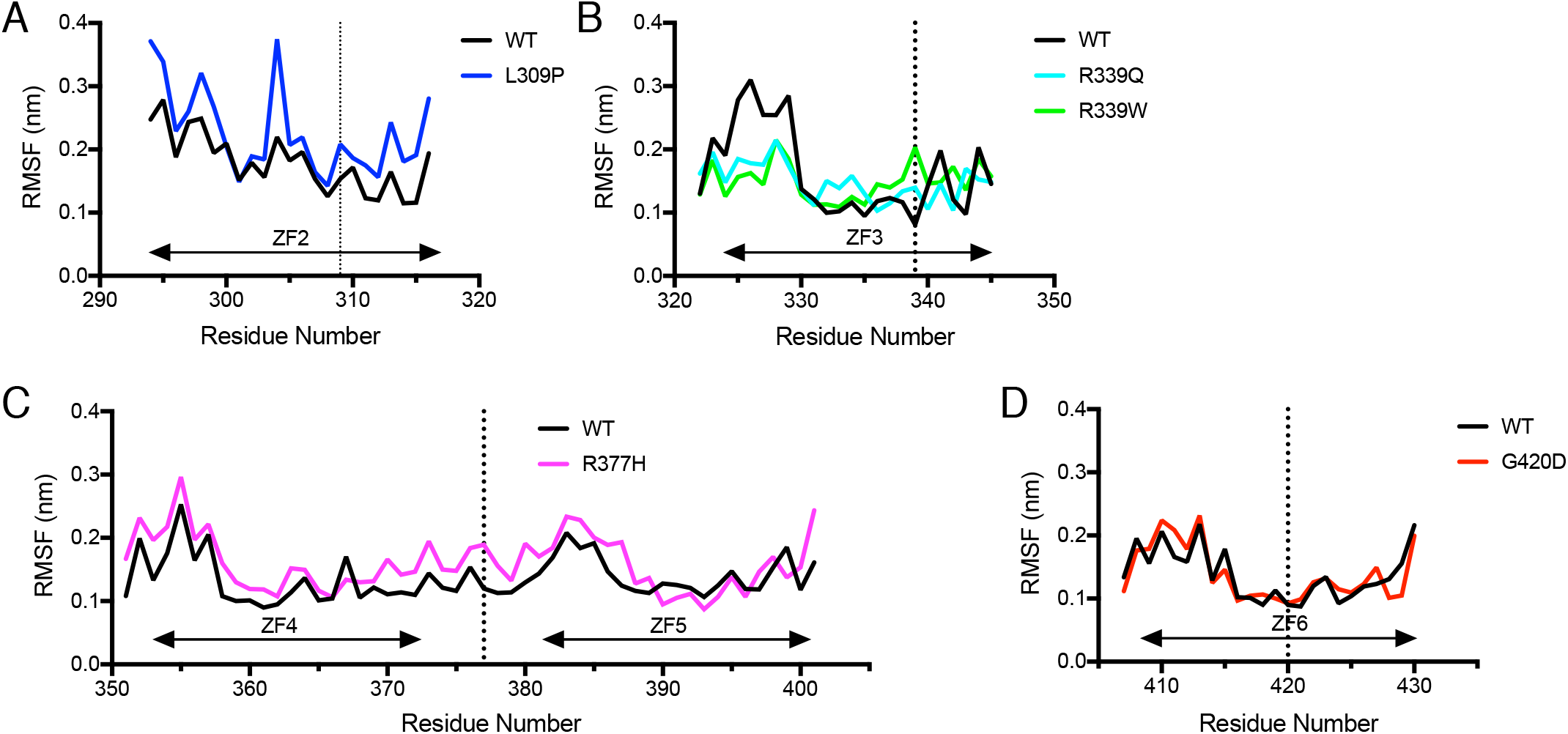
Flexibility of CTCF ZFs before and after mutation measured by MD simulations. **(A-D)** Backbone RMSF values during the MD simulations for mutant CTCF ZFs compared to WT, spanning their associated ZF domain; dotted vertical line indicates position of mutation. **(A)** L309P; ZF2 **(B)** R339Q & R339W; ZF3 **(C)** R377H; ZF4 & 5, **(D)** G420D; ZF6.

