## Supplementary Table 2 for "Somatic mutations in CTCF zinc fingers produce cellular phenotypes explained by structure-function relationships"

**Supplementary Table 2** Analysis of the distribution of missense somatic mutations and SNPs in CTCF. The number of expected mutations was determined from the proportion of mutations expected if they were evenly distributed between each domain. The observed/expected (O/E) ratio confirms if there is a de-enrichment (<1.0) or an enrichment (>1.0) of non-synonymous changes. Statistically significant differences are indicated in bold calculated using the Chi-square test.

**CTCF somatic missense mutations**

| *Domain* | *Observed* | *Expected* | *Observed/Expected Ratio* | *P* |
|---|---|---|---|---|
| N | 94 | 150 | *0.63* | ***P*<0.0001** |
| ZF | 266 | 181 | *1.47* | ***P*<0.0001** |
| C | 54 | 83 | *0.65* | ***P*=0.0067** |
| total | 414 | 414 | | |

**CTCF SNPs**

| *Domain* | *Observed* | *Expected* | *Observed/Expected Ratio* | *P* |
|---|---|---|---|---|
| N | 94 | 74 | *1.27* | ***P*=0.0399** |
| ZF | 42 | 88 | *0.48* | ***P*<0.0001** |
| C | 67 | 41 | *1.63* | ***P*=0.0032** |
| total | 203 | 203 | | |
